# Constitutive plasma membrane interaction of active Rho GEF Ect2 inhibits cortex contraction pulses

**DOI:** 10.64898/2026.06.03.729549

**Authors:** Kristina Mrug, Nina Schulze, Katarina Blazevic, Konstantina Feller, Johannes Koch, Leif Dehmelt, Perihan Nalbant

## Abstract

During cell division, multiple guanine nucleotide exchange factors (GEFs), including the mitotic regulator Ect2 and Lbc-type GEFs, regulate the dynamics of RhoA activity and actomyosin contractility. In interphase, Lbc-type GEFs are predominantly cytosolic and are transiently recruited to the plasma membrane via a Rho-dependent positive-feedback mechanism that generates stochastic pulses of Rho activity and cortical contractions. In contrast, during interphase, Ect2 is primarily sequestered in the nucleus away from the cytoplasm and the plasma membrane. Following nuclear envelope breakdown in mitosis, Ect2 is released into the cytosol and constitutively associates with the plasma membrane independently of Rho via a C-terminal polybasic cluster. Here, we mimicked this mitotic state by expressing a constitutively active Ect2 variant that is lacking its nuclear localization signal in interphase, to examine its interplay with Lbc-stimulated pulsatile Rho dynamics. We found that active cytosolic Ect2 suppressed pulsatile dynamics and promoted peripheral enrichment of Rho activity and myosin. Pulse suppression required Ect2 catalytic activity, Rho-dependent positive feedback, and constitutive plasma membrane association. In contrast, an Ect2 variant that was lacking constitutive plasma membrane association via the polybasic cluster, was not only incapable of pulse suppression but instead even stimulated strong pulsatile Rho dynamics. Together, our results identify constitutive plasma membrane association as a key distinction between Lbc-type GEFs and Ect2, and demonstrate that this property enables Ect2 to suppress pulsatile Rho activity dynamics, thereby promoting a stable, non-pulsatile contractile signaling regime during mitosis.

## Introduction

The precise spatio-temporal regulation of actomyosin contractility is integral for dynamic cell shape changes throughout the cell cycle. The small Rho GTPase RhoA, controls cell contractility via its effector ROCK (Rho associated kinase) which activates myosin by phosphorylation (Amano et al., 1996). In adherent interphase cells, Rho controls multiple types of dynamic contractile structures including the actin-myosin rich cell cortex and prominent actin stress fibers. During cell division, the actin-myosin cytoskeleton is reorganized in the entire cell cortex, first leading to the formation of the spheric mitotic cell shape with increased cortical stiffness (Stewart et al., 2011) followed by the robust localization of myosin to the equatorial cortex and the formation of the cytokinetic cleavage furrow (Green et al., 2012). Although multiple guanine nucleotide exchange factors (GEFs) that activate Rho and regulate spatio-temporal myosin contraction dynamics during mitosis have been identified, their individual roles and their functional interplay are not yet fully understood.

The RhoA GEF Ect2 (Epithelial Cell Transforming 2) is one of the best-studied mitotic Rho regulators and it was found to play a major role in controlling the formation of the contractile cleavage furrow during cytokinesis (Tatsumoto et al., 1999; Yüce et al., 2005). In interphase cells, Ect2 is localized in the nucleus and exists there in an autoinhibited activity state due to interactions of the catalytic GEF domain (Dbl-homology; DH) with the N-terminal BRCT domains and the pleckstrin homology (PH) domain (Kim et al., 2005; Saito et al., 2004b; Tatsumoto et al., 1999). Upon nuclear envelope breakdown, Ect2 becomes cytosolic where it promotes cortical Rho activation during early mitosis (Matthews et al., 2012). It has been suggested that cyclin dependent kinase 1 (Cdk1)-mediated phosphorylation during early mitosis only modestly enhances Ect2 GEF-activity (Niiya et al., 2006). During anaphase, when Cdk1 activity decreases, Ect2 becomes fully activated and is recruited to the spindle midzone through interaction with the Centralspindlin protein MgcRacGAP (Su et al., 2011a; Yüce et al., 2005). In addition, Ect2 constitutively interacts with the plasma membrane via its C-terminal PH and polybasic cluster (PBC) regions, an interaction that is particularly important at the equatorial membrane, where Ect2 is required for cleavage furrow ingression and successful cytokinesis (Kotýnková et al., 2016; Su et al., 2011a).

Several members of the Lbc-GEF family, including GEF-H1 and LARG, were also reported to regulate Rho activity and actomyosin contractility during cytokinesis and daughter cell abscission (Birkenfeld et al., 2007; Helms et al., 2016; Martz et al., 2013). In adherent interphase cells, Lbc-family RhoGEFs stimulate stochastic cortical Rho activity and actomyosin contraction pulses (Graessl et al., 2017; Kamps et al., 2020) which maintain the cortex as a dynamic and excitable network capable of rapid mechanical remodeling during dynamic cell shape changes (Bement et al., 2015). These subcellular pulses are generated via an activator-inhibitor signaling network coupling positive feedback between Lbc-family RhoGEFs and active Rho with delayed negative feedback mediated by actomyosin. Upon mitotic entry, the activity of these Lbc-family RhoGEFs is suppressed by mitotic kinase-dependent phosphorylation (Birkenfeld et al., 2007; Helms et al., 2016). During anaphase and cytokinesis, this inhibitory state is relieved, resulting in increased cortical Lbc-GEF signaling and contractility. Taken together, the activities of both Ect2 and Lbc-family RhoGEFs are reduced during early mitosis and subsequently increased during anaphase and cytokinesis, raising the question of how these regulators control spatio-temporal Rho activity and cell contraction dynamics during cell division.

Similar to other RhoGEFs, Ect2 and Lbc-family RhoGEFs activate Rho by stimulating the exchange of GDP for GTP on inactive Rho-GDP (Rossman et al., 2005; Tatsumoto et al., 1999). Interestingly, both Ect2 and Lbc-GEFs can also interact with active Rho, thereby promoting positive feedback amplification of Rho activity (Chen et al., 2020; Graessl et al., 2017). However, the mechanistic consequences of this interaction differ fundamentally. Lbc-GEFs are recruited to the plasma membrane by the interaction with active Rho, which closes a protein-localization-based positive feedback loop that is central to generate pulsatory Rho activity pulses (Graessl et al., 2017; Kamps et al., 2020). In contrast, Ect2 can localize to the plasma membrane independently of Rho via a polybasic cluster (PBC) in its C-terminus (Kotýnková et al., 2016; Su et al., 2011a). Binding to active Rho was shown to increase the enzymatic activity of Ect2 which thereby can mediate positive feedback amplification at the plasma membrane without an additional plasma membrane recruitment step (Chen et al., 2020). How these distinct positive feedback mechanisms might affect spatio-temporal Rho activity patterns, contraction dynamics and cell shape changes, and how they might interplay with each other, is currently not known.

Here, we found that constitutively active, plasma membrane associated Ect2 efficiently inhibits cortical Rho activity and myosin dynamics and instead stimulates a uniform, strong contractile state in the whole cell cortex. Loss of PBC-mediated constitutive plasma membrane association (ΔPBC) reverses this Ect2 function from inhibition to stimulation of local contraction pulses. Ect2ΔPBC-induced contraction pulses require binding to active Rho, suggesting that this mutant can mediate positive feedback via a mechanism that is similar to the previously described Lbc-GEF driven Rho signaling network. These results establish the PBC region as a critical module that couples membrane association to the suppression of pulsatile Rho activity dynamics, thereby enabling the function of Ect2 to stimulate a prolonged high contractile state in the cell cortex during mitosis.

## Results

In our previous studies, we found that Lbc-type GEFs can stimulate Rho activity pulses via positive feedback amplification of Rho activity. As Ect2 was recently also shown to be able to amplify Rho activity, we were interested, how this regulator affects Rho in cells. For these studies, we used the U2OS osteosarcoma cell model system previously established for our work on Lbc-type GEFs. As shown in Fig. 1A-C although wildtype Ect2 localized nearly exclusively to the nucleus, its expression still significantly stimulated the formation of contractile stress fibers. We next expressed an N-terminally truncated Ect2 variant that lacks the auto-inhibitory N-terminal region and the nuclear localization signal in the S-loop (ΔN-Ect2) (Saito et al., 2004a), thereby mimicking the active mitotic state of Ect2 following nuclear envelope breakdown. This constitutively active mutant strongly stimulated stress fiber formation and significantly elevated F-actin levels and cell rounding compared to either control (EGFP) or Ect2 wild-type (Fig. 1B-C). In contrast, expression of N-terminally truncated, catalytically inactive Ect2 variant (ΔN-Ect2-DHmut) did not induce these profound morphological changes (Fig. 1B-C).

**Figure 1:**
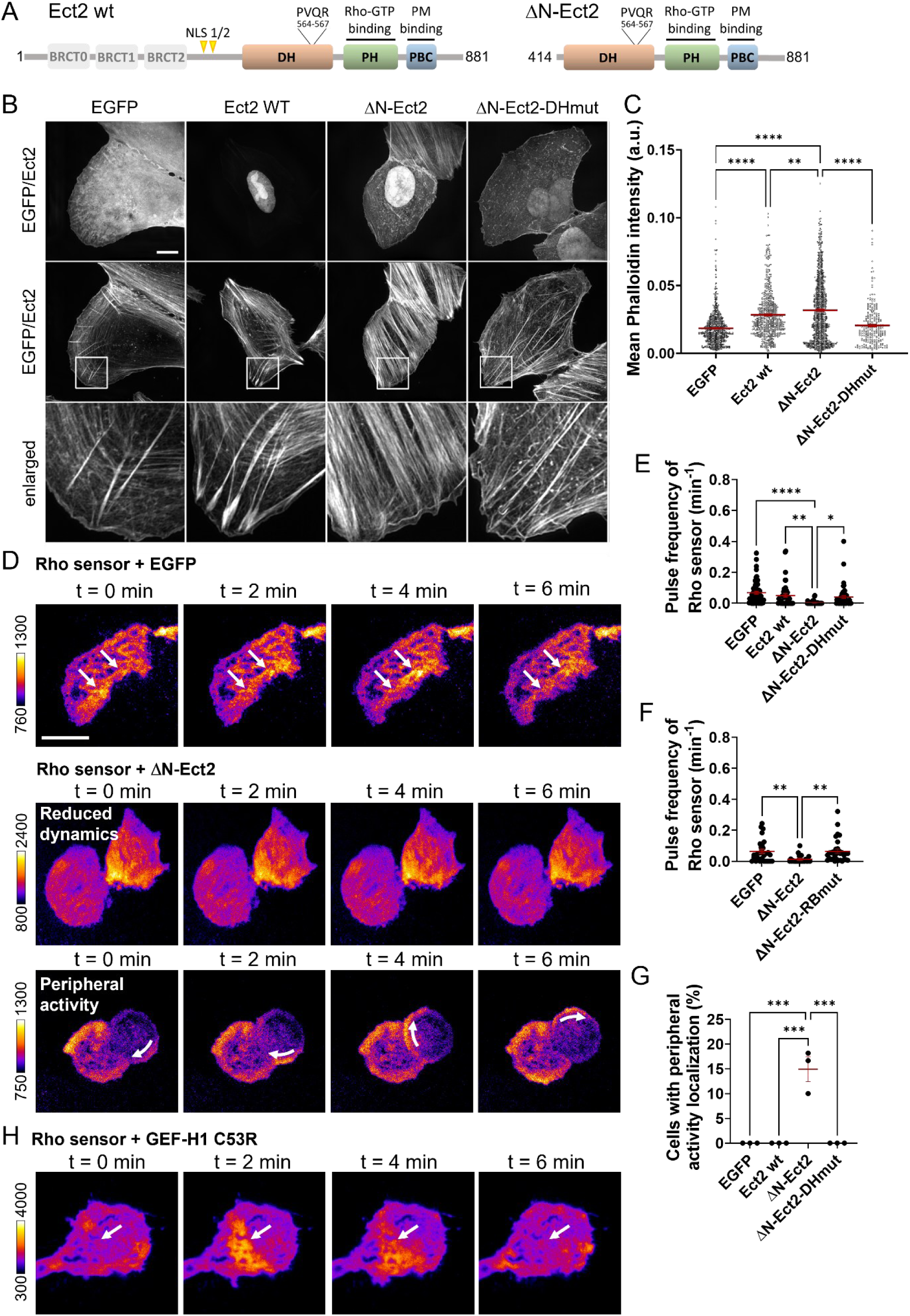
Active, plasma membrane-bound Ect2 suppresses spontaneous pulsatile Rho activity. **A)** Schematic for domain structures of Ect2 wild-type (Ect2-wt), N-terminally truncated active (ΔN-Ect2) and the catalytically inactive variant with PVQR->AAAA (564-567) substitutions (ΔN-Ect2-DHmut). **B)** Representative maximum intensity projections of 3D-SIM images of U2OS cells expressing EGFP control or the indicated EGFP-tagged Ect2 constructs. Upper panels show EGFP or EGFP-tagged Ect2 localization, middle panels display F-actin stained with phalloidin, and lower panels present magnified views of the regions depicted in the actin images. Scale bar: 10 µm. n = 7-15 cells. **C)** Quantification of mean phalloidin fluorescence intensity in wide-field images of cells expressing the indicated constructs. Cells were stained with WGA to visualize cell morphology and with phalloidin to label F-actin. Transfected cells were identified EGFP fluorescence. F-actin intensity was quantified by automated image analysis as described in the Methods. Data represent n ≥ 276 cells from 4 independent experiments. Statistical analysis was performed using one-way ANOVA followed by Tukey’s post hoc test. Bars indicate mean ± SEM. **D and H)** Representative TIRF images of the Rho activity sensor (mCherry-Rhotekin RBD) co-expressing either EGFP control, constitutively active Ect2 (EGFP-ΔN-Ect2) **(D)** or constitutively active GEF-H1 C53R **(H)**. Two phenotypes induced by active Ect2 are shown in **(D)**: (top) reduced pulsatory Rho sensor signal dynamics and (bottom) peripheral enrichment of Rho sensor signal with slow circumferential movement (white arrow). Frame rate: 3 frames/min, scale bar, 20 µm. **E and F)** Percent cells with peripheral Rho sensor enrichment **(E)** and the frequency of Rho sensor pulses in the central cell region **(F)**; n ≥ 32 cells from 4 independent experiments. **(G)** Average pulse frequency of the Rho activity sensor signal upon co-expression of EGFP-control, active EGFP-ΔN-Ect2 or a mutant which cannot bind to active Rho (EGFP-ΔN-Ect2-RBmut); n ≥ 25 cells from 3 independent experiments. Bars indicate mean ± SEM.

To measure how active Ect2 might affect Rho activity dynamics, we used a published variant (Nanda et al., 2023) of a translocation sensor based on the Rhotekin GTPase binding domain (GBD) (Benink and Bement, 2005). The sensor used here contains two tandem GBDs to increase affinity for active Rho and is expressed at very low levels due the delCMV promoter (Watanabe and Mitchison, 2002). Combined with TIRF microscopy, such sensors enable highly sensitive detection of sensor translocation from the cytosol to the plasma membrane upon binding to active Rho.

As shown in Fig. 1D-F, ectopic expression of active EGFP-ΔN-Ect2 strongly reduced spatio-temporal Rho activity dynamics in central cell attachment areas to a level which was barely detectable (Supplementary Video 1). This is in stark contrast to ectopic expression of the active Lbc-type GEF GEF-H1 C53R, which strongly stimulates such spatio-temporal Rho activity dynamics (Fig. 1H). Interestingly, a subset of ΔN-Ect2 transfected cells generated a slowly moving Rho activity wave that was exclusively enriched in peripheral attachment regions (Fig. 1D, G; Supplementary Video 1). The inhibitory effect of ΔN-Ect2 on stochastic pulsatory cortical Rho activity dynamics was observed already at relatively low Ect2 expression levels suggesting a potent inhibitory molecular mechanism (Supplementary Fig. 2). This effect of ΔN-Ect2 required catalytic GEF activity, as the corresponding inactive mutant (EGFP-ΔN-Ect2-DHmut) failed to inhibit stochastic cortical Rho activity dynamics and also did not induce slowly moving peripheral Rho activity waves (Fig. 1G). The effect of ΔN-Ect2 also required the interaction between Ect2 and active Rho as an Ect2 variant with mutations in the two amino acids that mediate this interaction (EGFP-ΔNEct2-RBmut (Chen et al., 2020)) did not alter Rho activity dynamics (Fig. 1F). Thus, binding to active Rho, and presumably the resulting positive-feedback amplification of Rho activity, is required for both the inhibitory effect of ΔN-Ect2 on stochastic pulsatory cortical Rho activity dynamics and the induction of slowly propagating peripheral Rho activity waves.

We next investigated if EGFP-ΔN-Ect2 can also suppress the strongly stimulated Rho activity dynamics induced by Lbc-GEFs. As previously shown (Chang et al., 2008), depolymerization of microtubules by addition of the pharmacological compound nocodazole can release and activate GEF-H1, and this leads to a strong increase in pulsatile cortical Rho activity dynamics (Fig. 2; Supplementary Video 2). In contrast, EGFP-ΔN-Ect2 completely blocked this nocodazole induced effect (Fig. 2; Supplementary Video 2). Thus, active Ect2 does not only efficiently inhibit basal stochastic Rho activity dynamics, but also suppresses the prominent pulsatile Rho activity dynamics that are strongly stimulated by Lbc-GEF activation via nocodazole.

**Figure 2:**
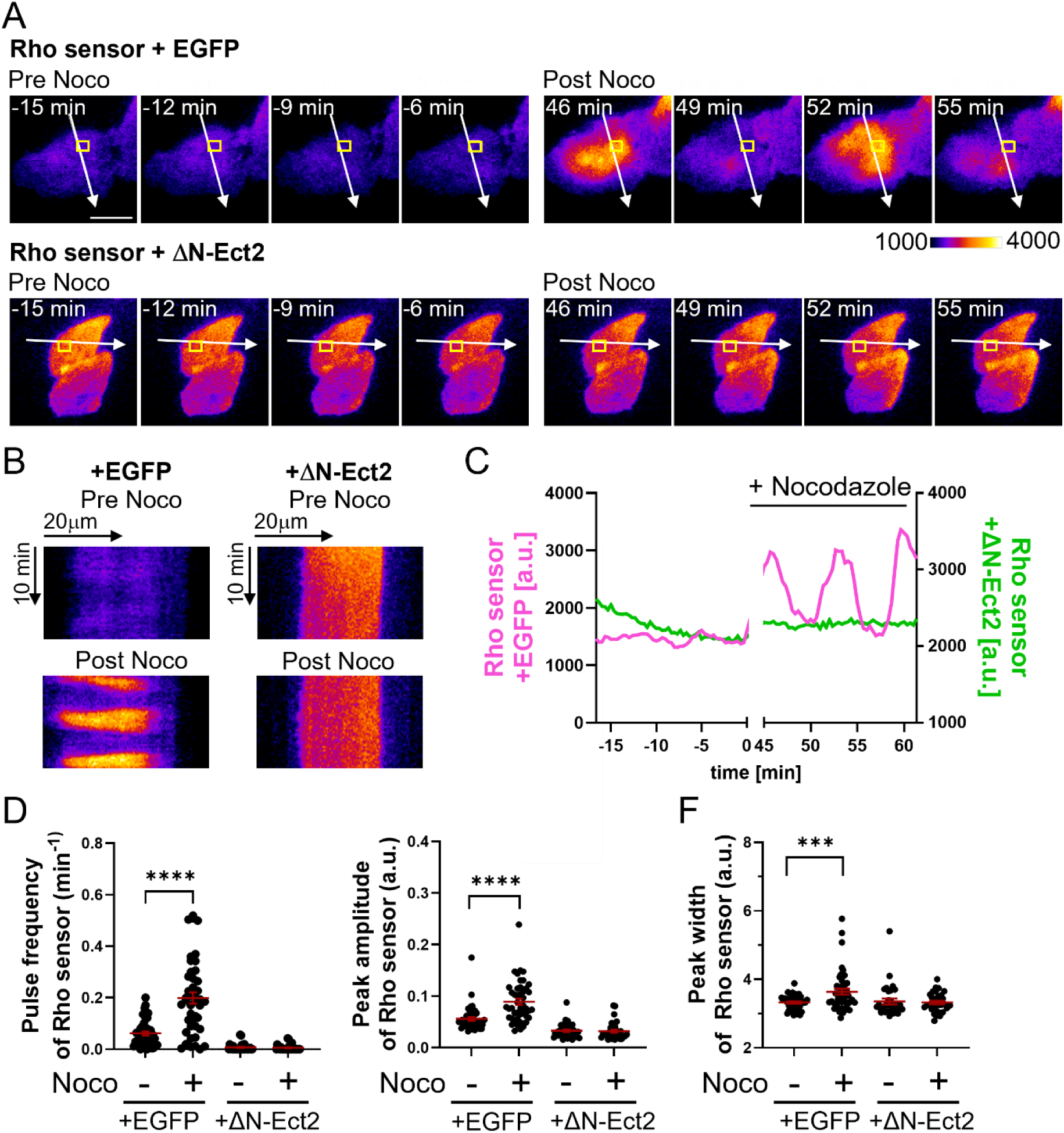
Active, plasma membrane-bound Ect2 suppresses Lbc-GEF stimulated pulsatile Rho activity. **A)** Representative TIRF time-lapse Rho sensor images of cells co-transfected with EGFP control (top panels) or EGFP-ΔN-Ect2 (bottom panels), prior (left) and post (right) nocodazole treatment). Frame rate: 3 frames/min, scale bar = 20 µm. **B)** Kymographs along the lines depicted in the cell images in **(A)**. **C)** Rho sensor signal intensity profiles in regions depicted in **(A)**. **(D-F)** Quantification of average Rho sensor pulse frequency **(D)**, peak amplitude **(E)** and peak width **(F)** pre- and post-nocodazole treatment. n ≥ 30 cells from 3 independent experiments. Statistical analysis was performed using one-way ANOVA followed by Tukey’s post hoc test. Bars indicate mean ± SEM.

Myosin II generates contractile actomyosin forces downstream of Rho (Amano et al., 1996). To investigate the functional consequence of ΔN-Ect2 expression, we generated a U2OS cell line stably expressing mCherry-Myosin-IIA and performed TIRF microscopy to monitor myosin-mediated contractility. Consistent with the observed changes in Rho activity, EGFP-ΔN-Ect2 markedly suppressed cortical myosin contraction dynamics in central regions of the cell and instead promoted the accumulation of myosin in peripheral cell attachment areas in a subset of cells (Fig. 3A-D). The peripheral myosin enrichment exhibited circumferential movement, similar to the Rho activity patterns observed in EGFP-ΔN-Ect2-expressing cells (Fig. 3A; Supplementary Video 3). This behavior was reflected in an increased standard deviation of myosin intensity over time (Fig. 3E, red data points). Using spinning disk confocal microscopy, we found that the circumferential movement was confined to adherent regions and did not extend along the apical-basal axis (Fig. 3F-G; Supplementary Video 4). This movement gave rise to periodic oscillations of the entire cell body, suggesting that ΔN-Ect2-mediated peripheral Rho activation induces strong anisotropic myosin-dependent contractile forces (Fig. 3G; Supplementary Video 4). Together, these data indicate that the ΔN-Ect2-induced transition in Rho activity from fast, stochastic pulses to spatially extended, slow peripheral waves is mirrored by a corresponding shift in myosin dynamics.

**Figure 3:**
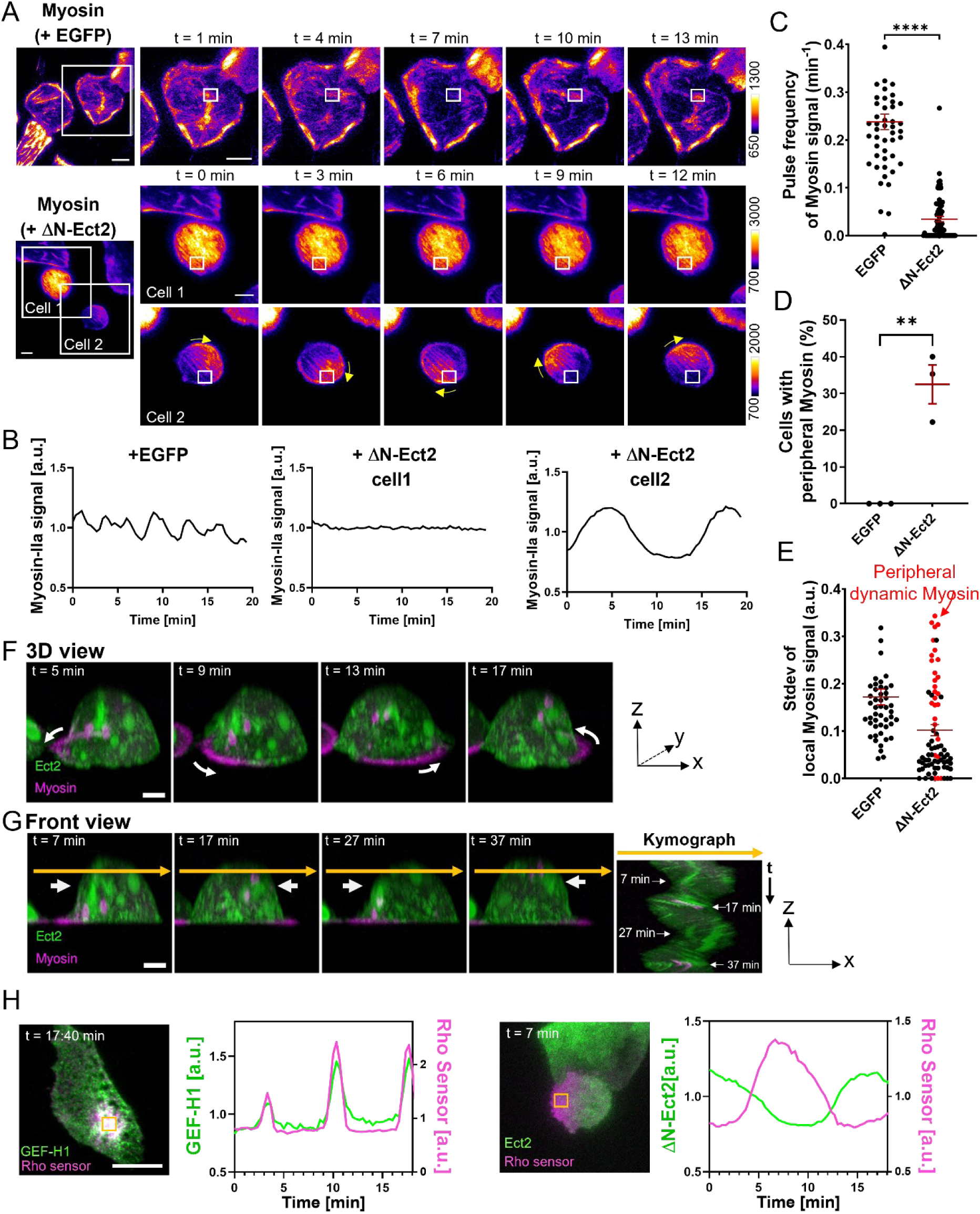
Active plasma-membrane bound Ect2 inhibits Myosin-IIA contraction pulses and generates strong peripheral Myosin-IIA enrichment. **(A)** Representative TIRF images of cells stably expressing mCherry-Myosin-IIA, transfected with either EGFP control or constitutively active Ect2 (EGFP-ΔN-Ect2). Two phenotypes induced by active Ect2 are shown: (cell1) reduced pulsatory Myosin-IIA signal dynamics and (cell2) peripheral enrichment of Myosin-IIA with slow circumferential movement (white arrow). Scale bars: 20 µm. **(B)** Mean normalized intensity values of the Myosin-IIA signal in regions marked in A (white boxes). **(C, D)** Average pulse frequency of Myosin-IIA signal **(C)** and percent cells with peripheral Myosin-IIA enrichment **(D)**. **(E)** Standard deviation of local Myosin-IIA signal in cells from **(C)** and **(D).** Cells with circumferential signal movement are depicted in red. n≥50 cells from 3 independent experiments. Unpaired t-test. **(F, G)** Representative 3D-projection of spinning disk confocal images of mCherry-Myosin-IIA (magenta) and active Ect2 (EGFP-ΔN-Ect2) (green). **(F)** 3D view (x, y, z): White arrows mark the movement of myosin signal maximum. **(G)** Front view (z, x): white arrows point to oscillatory movement of the cell body. The kymograph was generated along the orange arrow in front view cell images. n = 8 cells from 3 independent experiments. Scale bars: 5 µm. **(H)** Representative TIRF images of cells expressing the Rho activity sensor (magenta) together with constitutively active GEF-H1 (EGFP-GEF-H1 C53R, green; left) or constitutively active Ect2 (EGFP-ΔN-Ect2, green; right). Frame rates: 3 frames/min, scale bars = 20 µm. Bars indicate mean ± SEM.

Interestingly, we observed a significant delay between Ect2 localization and the peak of Rho activity of several minutes in these slow, peripheral waves (Fig. 3H; Supplementary Fig. 3; Supplementary Videos 5 and 6). In fact, Ect2 and Rho dynamics were nearly perfectly anti-phasic to each other. This is in stark contrast to Lbc-GEF stimulated Rho activity pulses in central cell attachment areas, in which the GEF and Rho are nearly perfectly in sync (Fig. 3H; Supplementary Video 5). While synchronous pulse dynamics observed for Lbc-GEFs and Rho are in agreement with a typical positive feedback mechanism, in which both molecules stimulate each other, the anti-phasic dynamics of active Ect2 and active Rho in slow, peripheral waves is highly counterintuitive and not compatible with mutual activation via localization. This suggests that the mechanism that underlies Lbc-GEFs stimulated Rho activity pulses and the mechanism that underlies Ect2-stimulated slow, peripheral Rho activity waves are fundamentally distinct.

This raises the question of why Lbc GEFs and Ect2 have such distinct effects on Rho activity dynamics. One key difference between these molecules is the mechanism by which they associate with the plasma membrane (Kotýnková et al., 2016). While the Lbc-GEFs GEF-H1 and LARG are predominantly cytosolic and are recruited to the plasma membrane through interactions with active membrane-bound Rho, Ect2 was reported to exhibit intrinsic plasma membrane affinity via its C-terminal polybasic cluster (PBC), and this PBC-dependent membrane localization was shown to be essential for cytokinesis (Kotýnková et al., 2016). Using spinning disk confocal microscopy, we confirmed that ΔN-Ect2 efficiently localized to the plasma membrane, and that this localization was strongly reduced in a mutant construct that lacked the PBC (ΔN-Ect2-ΔPBC) (Fig. 4A). Strikingly, in contrast to ΔN-Ect2, ΔN-Ect2-ΔPBC failed to suppress spatio-temporal Rho activity dynamics in central cell attachment regions, and it also did not induce the slow, peripheral Rho activity waves (Fig. 4B-F, Supplementary Video 7). These findings suggest that constitutive plasma membrane association of Ect2 is required to efficiently suppress pulsatile Rho activity dynamics.

**Figure 4:**
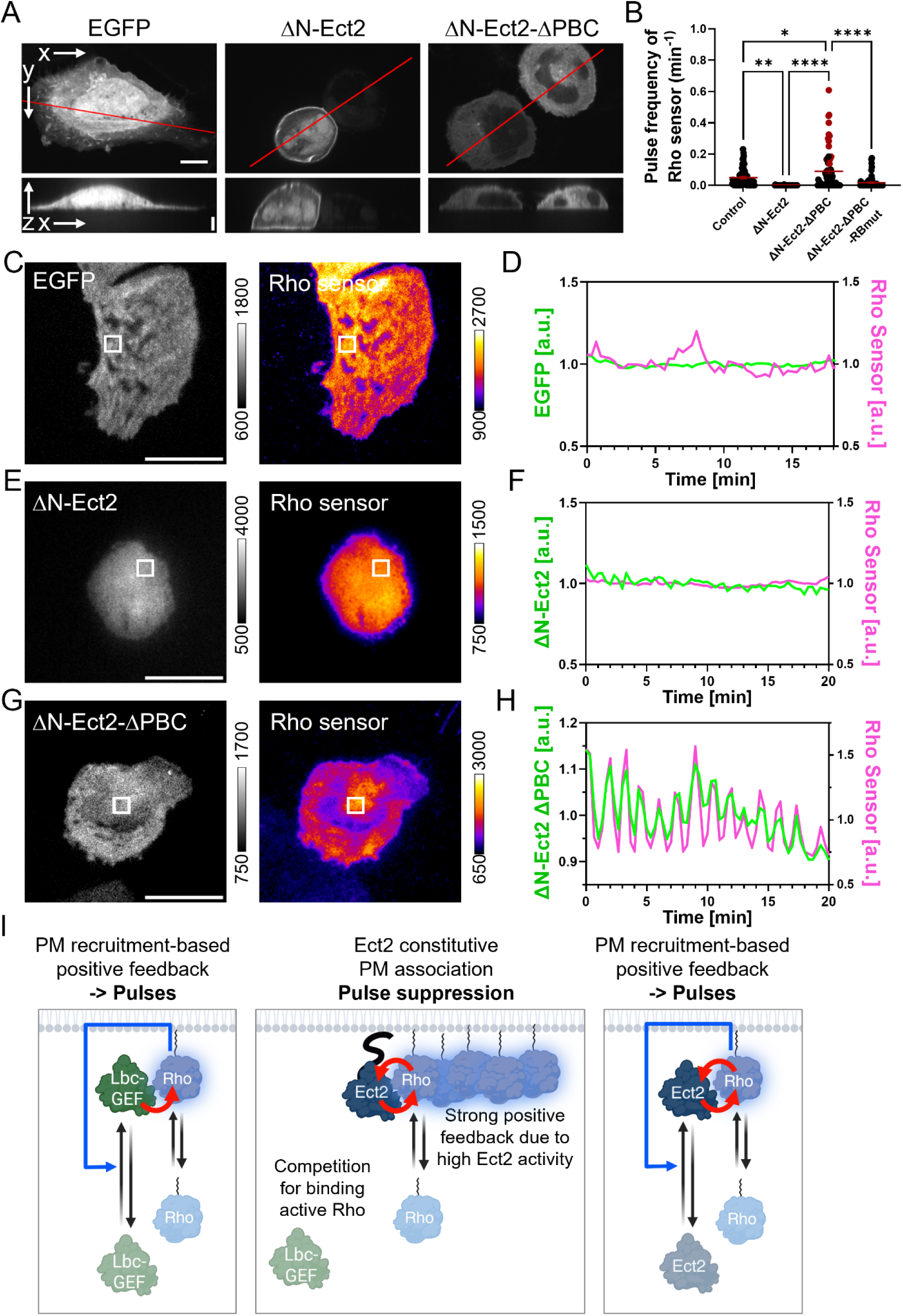
Plasma membrane binding deficient Ect2 stimulates Rho activity pulses. **A)** Representative spinning disk confocal images of fixed U2OS cells transfected with either EGFP control, active Ect2 (EGFP-ΔN-Ect2), or a variant lacking the C-terminal PBC region (EGFP-ΔN-Ect2-ΔPBC). A schematic of the constructs is shown in Supplementary Fig. 1A. Upper panels: single central z-plane, lower panels: side views show orthogonal (x-z) projections of the z-stack along the line indicated in the corresponding upper panels. Scale bars: 10 µm (xy), 5 µm (z), n=50-56 cells from 3 independent experiments. **B)** Average pulse frequency of the Rho activity sensor signal (mCherry-Rhotekin-RBD) in cells expressing the indicated Ect2 variants. Red dots mark cells that generate fast high-amplitude pulses. n ≥ 51 cells from 3 independent experiments. using one-way ANOVA followed by Tukey’s post hoc test. Bars indicate mean ± SEM. **C, E, G)** Representative TIRF images of cells expressing the Rho activity sensor (right) and the indicated constructs (left), respectively. Frame rate: 3 frames/min, scale bar = 20 µm. **D, F, H)** Mean normalized intensity values of the Rho sensor signal in the corresponding cell regions (white boxes) in **(C, E, F)**. **I)** Proposed interplay between Ect2 and pulsatory Rho contraction signal network dynamics. Left: Pulsatory Rho contraction depends on a positive feedback loop in which Lbc-GEFs are recruited to the plasma membrane through binding to active Rho (right angled blue arrow). Curved red arrow points to enzymatic activation. Middle: In contrast, Ect2 is constitutively associated with the plasma membrane, independently of active Rho (Su et al., 2011b). Increased local concentration of Ect2 may result in a higher effective on-rate, thereby conferring a kinetic advantage that can outcompete Lbc-GEFs and thereby suppress Rho activity pulses. Curved red arrows illustrate enzymatic activation and allosteric activation. Right: Loss of the C-terminal PBC reduces Ect2 plasma membrane association, shifting its properties towards a Lbc-GEF-like phenotype, that is based on plasma membrane recruitment to active Rho, and that stimulates Rho activity pulses.

Instead, ΔN-Ect2-ΔPBC already by itself was able to induce high-frequency, high-amplitude Rho activity pulses in a substantial subset of cells (Fig. 4B, red data points; representative example shown in Fig. 4G-H). This phenotype suggests that Ect2 lacking constitutive PBC-mediated plasma membrane association promotes pulsatile Rho activity dynamics, reminiscent of those generated by Lbc-family RhoGEFs (Graessl et al., 2017). Consistent with the localization-based positive feedback mechanism reported for Lbc-type GEFs, ΔN-Ect2-ΔPBC and Rho activity dynamics were nearly perfectly in sync, suggesting that these molecules mutually stimulate each other via plasma membrane recruitment (Fig. 4H, I). Indeed, the ability of Ect2 to bind active Rho is critical for this effect, as cytosolic Ect2 lacking binding to active Rho (ΔN-Ect2-ΔPBC-ΔRB) failed to induce high-frequency Rho activity pulses (Fig. 4B).

Together, these results suggest that active, cytosolic Ect2 can drive highly dynamic, pulsatile Rho activity. Conversely, constitutive Ect2 association with the plasma membrane suppresses these dynamics.

## Discussion

Here, we show that an active variant of the mitotic Rho GEF Ect2 can efficiently suppress Rho activity and contraction pulses that are stimulated by Lbc-type GEFs. In our studies, we used an established truncated Ect2 variant (ΔN-Ect2) that lacks the N-terminal auto-inhibitory BRCT domains and the S-loop, resulting in strongly increased activity and predominantly cytosolic localization. This active Ect2 not only suppressed basal stochastic pulses in Rho activity but also inhibited high-amplitude pulses and waves of Rho activity and myosin contraction induced by activation of the Lbc-GEF GEF-H1 upon nocodazole treatment. These findings highlight the robustness of this mechanism, as it suppresses even pronounced pulses and waves of Rho activity driven by strong Lbc-GEF activation.

Neither a catalytically inactive DH mutant nor wild-type Ect2 elicited these effects, demonstrating that basal Ect2 activity is insufficient and that increased GEF activity is required to suppress Rho activity and myosin. In addition, suppression of pulsatory Rho dynamics by Ect2 also depends on its interaction with active Rho. A mutant defective in the Rho-binding region failed to suppress Rho activity pulses, consistent with biochemical evidence that active Rho binding enhances Ect2 catalytic activity. These findings suggest that Rho-dependent activation of Ect2 is required to drive Ect2 into the high-activity state necessary to suppress pulsatile Rho signaling.

Lbc-GEF-mediated pulsatory Rho dynamics are driven by a positive feedback loop in which active Rho recruits Lbc-GEFs to the plasma membrane, thereby promoting further Rho activation (Graessl et al., 2017; Kamps et al., 2020). The observation that active Ect2 suppresses even strong GEF-H1-induced pulses upon nocodazole treatment indicates that the Ect2-dependent inhibitory mechanism can override this pulsatory circuit. One possible explanation is that Ect2 competes with Lbc-GEFs for binding to active Rho, thereby limiting Lbc-GEF membrane recruitment and disrupting its localization-based positive feedback loop.

In a subpopulation of cells, active Ect2 expression led to the accumulation of Rho activity and myosin at the cell periphery, where they formed contractile signals that propagated slowly in a circumferential manner. This behavior differs from previously reported traveling waves of Rho activity in dividing *Xenopus* and starfish eggs (Bement et al., 2015; Michaud et al., 2022), as it appears to be confined to sites of cell adhesion and does not extend into the three-dimensional cell cortex.

In the slowly propagating activity waves induced by active Ect2, the temporal relationship between Rho and myosin signals is in agreement with established cell contraction signalling. Consistent with previous reports (Graessl et al., 2017; Kamps et al., 2020), the myosin signal maximum followed Rho activity with a measurable delay. However, the temporal relationship between active Ect2 plasma membrane localization and the peak of Rho activity was very unusual, with a prolonged delay, indicating that Ect2 accumulation does not translate directly into local enrichment of active Rho. Notably, this delay spans several minutes, leading to nearly anti-phasic dynamics and thus contrasts strongly with the almost synchronous, in-phase dynamics measured for positive feedback between Lbc-type GEFs and Rho. This marked temporal separation therefore suggests that a different regulatory mechanism controls these dynamics, which might include additional regulators or spatial constraints.

In addition to the canonical tandem DH-PH module shared with other Dbl-family GEFs, the truncated active Ect2 variant contains a C-terminal polybasic cluster (PBC) that mediates constitutive association with the plasma membrane and is essential for successful cytokinesis (Kotýnková et al., 2016; Su et al., 2011a). This mode of membrane association differs from that of Lbc-family GEFs, which rely on interaction with active Rho for membrane recruitment as part of the positive feedback loop underlying pulsatile Rho contractility (Graessl et al., 2017). Our data suggest that the presence of the PBC may define two functionally distinct states of Ect2: a plasma membrane-associated state that suppresses pulsatory contraction signals, and a second state that stimulates pulse generation.

Previous studies have shown that phosphorylation of the PBC region by Cdk1 prevents Ect2 association with the plasma membrane until anaphase, after which its localization is restricted to the equatorial region by centralspindlin (Su et al., 2011a). In agreement with this phosphorylation-mediated control of the PBC-dependent membrane anchor, only a subpopulation of cells exhibited pulsatile contraction dynamics upon expression of the active Ect2 construct lacking the PBC. This heterogeneity may thus reflect cell-to-cell variability in regulatory states. As the experiments were performed in non-synchronized interphase cells, differences in phosphorylation status, for example due to elevated mitotic kinase activity in a subset of cells, could influence Ect2 function. Similar to the pulsatory Rho contraction signaling network driven by Lbc-family GEFs (Graessl et al., 2017), Ect2-mediated contraction pulses also required binding to active Rho. This suggests that positive feedback is a core driver of the Ect2-dependent pulsatile signaling network.

Together, these observations suggest that, in the absence of the PBC, Ect2 adopts a pulse-generating state driven by self-organizing signaling dynamics. In contrast, PBC-mediated membrane association appears to suppress the emergence of such pulsatile behavior, suggesting that constitutive membrane anchoring constrains the spatial or dynamic coupling required for feedback-driven amplification. Notably, this inhibitory effect is not limited to Ect2-driven dynamics, as Ect2 expression also suppressed strong Rho activity pulses and waves induced by activation of the Lbc-family GEF GEF-H1 upon nocodazole treatment, indicating a broader interference with pulsatile Rho dynamics. This effect may arise from the increased effective local concentration of membrane-associated Ect2, enabling it to outcompete cytosolic Lbc-GEFs and thereby override pulsatory dynamics (Fig. 4I).

The observations raise the question, why Ect2 and Lbc-GEFs are activated at the same time in late stages of mitosis. We hypothesize that the combined action of these regulators may be particularly relevant during cytokinesis, where multiple morphological processes must be coordinated simultaneously. At this stage, cells must establish a mechanically robust and stable cleavage furrow, and at the same time initiate daughter cell adhesion and cell separation. Thus, the coordinated activity of Ect2 and Lbc-GEFs may contribute to balancing stable contractility with dynamic spatiotemporal regulation of actomyosin activity. As Ect2 and members of the Lbc-family of Rho GEFs are activated in parallel, they can contribute to these mitotic and migratory behaviors. Coordination between these GEF activities is thus required to spatiotemporally organize the contrasting contractile regimes, balancing a stable, mechanically robust cleavage furrow with the more dynamic and less stable cortical contractions that enable daughter cell separation.

## Author Contributions

P.N. conceived the study, supervised the project, and wrote the manuscript. K.M. performed and analyzed the majority of experiments. K.B. generated the stable cell line and conducted parallel live-cell imaging of Ect2, Rho activity, and Myosin-IIA dynamics in individual cells. K.F. performed imaging and quantitative analysis of pulsatile Myosin-IIA dynamics following overexpression of Ect2 variants. N.S. and J.K. carried out 3D-SIM imaging. N.S. additionally performed 3D spinning-disk confocal microscopy of fixed and live cells, as well as epifluorescence imaging and analysis of cell morphology and actin intensity. L.D. contributed to image and data analysis, and optimization. All authors contributed to data interpretation, discussion and figure preparation.

## Acknowledgements

We acknowledge the use of the imaging equipment and the support in microscope usage by the “Imaging Center Campus Essen” (ICCE), Center of Medical Biotechnology (ZMB), University of Duisburg-Essen. This work was funded by the Deutsche Forschungsgemeinschaft (DFG, German Research Foundation) through CRC 1430 (Project ID 424228829; awarded to N.S. and P.N.) and by the DFG Heisenberg Program (grants DE 823/6-1 and DE 823/8-1) as well as the Principal Investigator Grant DE 823/10-1 to L.D.. The Andor/Nikon Spinning Disk confocal microscope was funded by the DFG-Project-ID 181168673. The Nikon Ti2-E TIRF DualCam microscope was funded by the DFG-Project-ID 361032973. The Nikon N-SIM S microscope was funded by the DFG–Project-ID 496847469.

## Materials and methods

### Cell culture, transfections and staining

Human U2OS osteosarcoma cells (HTB-96; ATCC) were maintained at 37 °C and 5 % CO_2_ humidified atmosphere using standard cell culture techniques (DMEM + GlutaMAX™ medium, 10 % FBS, Life technologies; Gibco). Imaging medium contained DMEM w/o Phenol red (Life Technologies), 10 % FBS, 10 mM HEPES, 1 mM CaCl_2_ and 1 mM MgCl_2_. To stimulate Rho activity via GEF-H1 release from microtubules cells were treated with the microtubule-disrupting agent nocodazole (30 µM, 45 min, Sigma Aldrich).

For imaging, cells were prepared by seeding onto either glass-bottom dishes (P35G-1.5-14-C, MatTek), µ-slide 8 well Glass Bottom chambers (ibidi, 80827) or glass coverslips (No. 1.5; VWR, MENZCB00130RAC). All glass surfaces were pre-coated with collagen type I (0.01 mg/ml, C8919-20ML, Merck) for 1 h prior to cell seeding. Transfection of plasmid DNA was performed using Lipofectamine 2000 (Life Technologies; Gibco). The number of cells, quantity of DNA and amount of Lipofectamine used are specified in the corresponding methods section.

For fluorescent staining, cells were fixed 18–24 h after transfection with 4% paraformaldehyde (PFA) for 20 min at 37 °C, followed by washing with DPBS. For plasma membrane labeling, cells were incubated with Wheat Germ Agglutinin (WGA) Alexa Fluor 647 conjugate (Thermo Fisher Scientific, W32466, 1:200 in HBSS, 10 min at RT) prior to permeabilization. Cells were subsequently permeabilized with 0.2% Triton X-100 in DPBS for 10 min at RT, washed with DPBS, and stained with rhodamine-phalloidin (Thermo Fisher Scientific, R415, 1:1000 in 2% BSA in DPBS, 1 h at RT) to visualize F-Actin and with DAPI (Thermo Fisher Scientific, D3571, 0.5µg/ml in DPBS, 10 min at RT) for nuclear staining. Samples were mounted using either ibidi Mounting Medium (ibidi, 50001) or ProLong Diamond Antifade Mountant (for 3D SIM, Thermo Fisher Scientific, P36961).

### Plasmid constructs

The Rho activity sensors delCMVmCherry-RBD and delCMVmCitrine-2xRBD were previously described (Graessl et al., 2017; Nanda et al., 2023)(Graessl et al., 2017 and Nanda et al.). pEGFP-Ect2 WT und pEGFP-ΔN-Ect2 (pEGFP-Ect2 414-882; ΔN5: constitutive active as described in (Saito et al., 2004a) and the catalytically inactive variant carrying the PVQR->AAAA substitution at residues 564-567 (pEGFP-ΔN-Ect2-DHmut) were provided by J. Birkenfeld (BioCopy AG, Basel, Switzerland). pEGFP-C1 (Clontech) was provided by L. Dehmelt (TU Dortmund University, Germany).

The plasmid encoding truncated, constitutively active Ect2 that lacks binding to active Rho (pEGFP-ΔN-Ect2-RBmut (Chen et al., 2020)) was generated by two sequential rounds of PCR-based site-directed mutagenesis followed by Gibson assembly using KpnI and AgeI sites. In the first round, the F621A substitution was introduced into pEGFP-ΔN-Ect2 using the following primers (Eurofins): 5′-CAAATTGCTGATGTTGTTGCTGAAGTAGATGGATGCCC-3′ (forward) and 5′-GGGCATCCATCTACTTCAGCAACAACATCAGCAATTTG-3′ (reverse). In the second round, the Y625A substitution was introduced on the F621A background to generate the double mutant (F621A/Y625A) using the primers: 5′-CAAATTTTTGATGTTGTTGCTGAAGTAGATGGATGCCC-3′ (forward) and 5′-GGGCATCCATCTACTTCAGCAACAACATCAAAAATTTG-3′ (reverse).

To generate the constitutively active Ect2 variant lacking the C-terminus (pEGFP-ΔN-Ect2-ΔPBC), the fragment encoding amino acids 414-766 was amplified from pEGFP-ΔN-Ect2 by PCR using the following primers (Eurofins): 5′-CGTCAGATCCGCTAGCGCTAGCCACCATGGTGAGCAAG-3′ (forward) and 5′-GGATCCCGGGCCCGCGGTACTTATTTACAAATGGTGTTAGCTACATGTC-3′ (reverse). The resulting PCR product was inserted into the vector backbone by Gibson assembly at the KpnI and AgeI restriction sites. To abolish binding of this variant to active Rho (pEGFP-ΔN-Ect2-ΔPBC-RBmut (Chen et al., 2020)), two point mutations (F621A and Y625A) were introduced by PCR-based mutagenesis, followed by Gibson assembly as described for pEGFP-ΔN-Ect2-RBmut.

The ΔN-Ect2-miRFP construct was generated by PCR-mutagenesis (primer: 5’-GGATCACCGCGCTTGAGAGCTCCGGACTCAGATCTCGAG-3’; 5’-ATAAACAAGTTAACAACAACCTATATCAAATGAGTTGTAGATCTACTTAACG-3’, Eurofins). The resulting PCR product was inserted into p-miRFP670-N1 (Addgene plasmid #79987) using Mfel and BlpI sites via Gibson assembly.

### Generation of stable cell line expressing mCherry-MyosinIIa

Cargo DNA was first cloned into a piggyBac transposon vector. To this end, the puromycin resistance cassette in pPBCAG-cHA-IRES-Puro (kindly provided by Christian Schröter, Max-Planck-Institute of Molecular Physiology, Dortmund) was replaced with a hygromycin resistance gene derived from pBabe-Hygro (Addgene plasmid #1765). The hygromycin cassette was amplified by PCR and inserted via Gibson assembly using the ClaI and MscI restriction sites. mCherry-tagged Myosin IIa was PCR-amplified from pCMV-mCherry-NMHCIIA (Dulyaninova et al., 2007; Addgene plasmid #35687) using the following primers: 5′-TCATTTTGGCAAAGAATTCCGCCACCATGGTGAGCAAG-3′ and 5′-CGATATCAAGCTTATCGAGCTTATTCGGCAGGTTTGGC-3′. The resulting fragment was cloned into the modified pPBCAG_cHA_IRES_Hygro vector using NotI and XhoI restriction sites. Insertion of the mCherry-Myosin IIa fragment into the vector disrupted and thereby eliminated the cHA multiple cloning site. Stable genomic integration was achieved in two steps. First, pPBCAG_mCherry-Myosin-IIa_IRES_Hygro was co-transfected with a piggyBac transposase expression vector (kindly provided by Christian Schröter) at a 2:1 ratio. After 48 h, antibiotic selection was initiated and maintained for two weeks. Cells were subsequently cultured in medium supplemented with 150 µg/mL hygromycin B (Invitrogen).

### Microscopy

#### Automated Widefield Fluorescence Microscopy

For quantification of cell morphology and F-actin intensity, 2x10^4^ cells were seeded onto µ-Slide 8 Well Glass Bottom chambers. The following day, cells were transiently transfected with 50 ng DNA and 0,8 µl Lipofectamine2000. Fluorescent staining was performed as described above using WGA Alexa Fluor 647, rhodamine-phalloidin, and DAPI. Samples were mounted using ibidi Mounting Medium. Imaging was performed using an automated EVOS M7000 imaging system (Thermo Fisher Scientific) equipped with a monochrome CMOS camera and an Olympus UPLXAPO 20×/0.8 NA objective (AMEP4906). Images were acquired using LED-based widefield epifluorescence illumination with sequential multi-channel acquisition. The acquisition protocol included DAPI (357/44 nm excitation, 447/60 nm emission), GFP (482/25 nm excitation, 524/24 nm emission), RFP (542/20 nm excitation, 593/40 nm emission), and Cy5 (628/40 nm excitation, 692/40 nm emission) fluorescence channels using predefined EVOS light cube configurations. Images were recorded in monochrome mode without binning. Channel-specific exposure times, gain settings, and LED illumination intensities were optimized prior to acquisition and subsequently maintained constant across all experimental conditions. Acquisition parameters were as follows: DAPI, 30 ms exposure time, gain 1, LED intensity 10%; GFP, 30 ms exposure time, gain 10, LED intensity 2%; RFP, 30 ms exposure time, gain 2, LED intensity 6%; and Cy5, 50 ms exposure time, gain 5, LED intensity 10%. For each condition, 60 fields of view were acquired at a single focal plane in a 6 × 10 grid using a serpentine horizontal stage-scanning pattern combined with a spiral inward acquisition sequence. Autofocus was performed in the DAPI channel using a fluorescence-optimized contrast-based autofocus algorithm. All images were acquired as monochrome images with 12-bit dynamic range and exported as uncompressed TIFF files.

#### Total Internal Reflection Fluorescence (TIRF) Microscopy

For quantification of subcellular activity dynamics, 1x10^5^ cells were seeded onto 35 mm Dishes. The following day, cells were transiently transfected with 300 ng DNA and 3 µl Lipofectamine2000. Prior to the imaging 18–24 h later, the medium was exchanged for imaging medium. TIRF microscopy was performed using a Nikon Ti2-E TIRF DualCam system equipped with an H-TIRF module for automated TIRF angle adjustments and an Andor Technology iXon Life DU-888 back-illuminated EMCCD camera. Excitation was provided by a diode laser cw LuXX 488 nm (250 mW) and a DPSS laser cw Jive 561 nm (300 mW) coupled through an Omicron LightHUB-4 compact laser beam combiner with AOTF-based laser modulation. Fluorescence excitation and emission were separated using an NSTORM QUAD dichroic mirror (reflection bands: 497–553 nm and 575–628 nm). Emission signals were collected using Semrock single band emission filters F37-516 for EGFP detection (525/50 nm) and F37-609 for RFP detection (600/50 nm). Cells were imaged using a Nikon CFI Apochromat TIRF 60×/1.49 NA oil immersion objective. Image acquisition was controlled using NIS-Elements Advanced Research (AR) software. Acquisition parameters were as follows: 488 nm: 100 ms exposure time, 10 % laser power; 651 nm: 200 ms exposure time, 10 % laser power. Live-cell imaging experiments were performed in a temperature-controlled incubation chamber maintained at 37 °C using imaging medium. Time-lapse image sequences were acquired at the indicated frame rates.

#### Spinning-disk Confocal Microscopy

For analysis of Myosin-IIa and dN-Ect2 dynamics in whole cell volumes, cells were prepared as described for TIRF microscopy. Live-cell imaging was performed on an inverted Nikon Ti-E microscope (Nikon Europe B.V., Amstelveen, The Netherlands) equipped with a Yokogawa CSU-X1 spinning-disk confocal unit (Yokogawa Electric Corporation, Tokyo, Japan) and controlled using Andor iQ software (version 3.7.0; Andor Technology, Belfast, UK). Images were acquired using an Andor Zyla 4.2 sCMOS camera (Andor Technology, Belfast, UK) in 16-bit mode with 2×2 binning. Cells were imaged using a Nikon CFI Apo TIRF 60×/1.49 NA oil immersion objective (Nikon Europe B.V., Amstelveen, The Netherlands). Excitation was provided by an Andor ILE multimode laser engine using 488 nm and 561 nm laser lines (Andor Technology, Belfast, UK). Emission was detected using Semrock BrightLine filters FF01-512/630-25 for 488 nm imaging and FF01-593/40-25 for 561 nm imaging (Semrock, Rochester, NY, USA). Acquisition parameters were as follows: 488 nm channel: 25 ms, 20% laser intensity; 561 nm channel: 50 ms exposure time, 30% laser intensity. Z-stacks were acquired with a constant z-step spacing of 0.4 µm and adjusted to cover the entire cellular volume. Time-lapse image series were acquired every 20 s for a total duration of 40 min (120 time points). Live-cell imaging was performed at 37 °C in a temperature-controlled incubation chamber manufactured by EMBLEM (Heidelberg, Germany).

For analysis of plasma membrane localization, 2x10^4^ cells were seeded onto µ-Slide 8 Well Glass Bottom chambers and transiently transfected with 50 ng DNA and 0,8 µl Lipofectamine 2000. Following fixation 18–24 h after transfection, samples were mounted using ibidi Mounting Medium. Fixed-cell imaging was performed on the same microscope setup as described for live-cell imaging, except, that images were acquired using 1×1 binning at a resolution of 1024×1024 pixels. The 488 nm channel was acquired with an exposure time of 200 ms and laser intensity setting of 8%. Z-stacks were acquired with a constant z-step spacing of 0.22 µm covering the entire cellular volume.

#### Structured Illumination Microscopy (SIM)

For visualization of the actin morphology by 3D-SIM, 1x10^5^ cells were seeded onto glass coverslips. Cells were transiently transfected with 300 ng plasmid DNA using 3 µl Lipofectamine 2000. Transfected cells were fixed, stained for F-actin and mounted with ProLong Diamond Antifade. 3D-SIM imaging was performed using an N-SIM S microscope (Nikon Europe B.V., Amstelveen, Netherlands) equipped with an ORCA-Fusion BT sCMOS camera (Hamamatsu Photonics K.K., Hamamatsu, Japan) and a CFI SR HP Apochromat TIRF 100×/1.49 NA oil immersion objective. Image acquisition was controlled using NIS-Elements AR software (version 5.42.07; Nikon). Excitation was provided by a ZIVA solid-state light engine (Lumencor, Beaverton, OR, USA). Fluorescence emission was collected through a quad-band dichroic mirror (405/488/561/640 nm) and detected with single-band emission filters. EGFP 476 nm excitation (25% laser power; 100 ms exposure time) with a SIM 488 BA emission filter (500–545 nm). Rhodamine-phalloidin was imaged using 545 nm excitation (20% laser power; 50 ms exposure time) with a SIM 561 BA emission filter (570–640 nm). Images were acquired at 2048×2048-pixel resolution using 16-bit image depth and 1×1 camera binning. For each field of view, complete cellular volumes were acquired as z-stacks with a step size of 0.2 µm. Raw SIM datasets were reconstructed in NIS-Elements using the standard stack reconstruction algorithm and default reconstruction parameters. Maximum intensity projections of reconstructed image stacks are shown. Image data storage and downstream processing were performed using OMERO (version 5.29.1).

### Image processing and data analysis

Microscopy image panels were generated using ImageJ/Fiji Software (https://imagej.net/software/fiji/) (Schindelin et al., 2012) and OMERO (version 5.29.1; https://www.openmicroscopy.org/omero/) (Allan et al., 2012). The image stabilizer plugin (K. Li, “The image stabilizer plugin for ImageJ,” http://www.cs.cmu.edu/~kangli/code/Image_Stabilizer.html, February 2008) was used to correct lateral drift. Representative images were adjusted with respect to brightness levels, cropping, scaling, and false color-coding via look-up tables. All images corresponding to the compared conditions were processed identically. To quantify pulsatile Rho activity and myosin-based contraction dynamics, a custom-made ImageJ script which was described previously (Graessl et al., 2017) was used. Statistical analyses and data plotting were carried out using Prism 10.3 (GraphPad). The type of statistical tests and significance are indicated in the respective figure legends. P values are indicated in the figures (ns, P > 0.05; *, P < 0.05; **, P < 0.01; ***, P < 0.001; ****, P < 0.0001).

#### Quantitative cell morphology and F-actin density analysis

Widefield fluorescence images were preprocessed in Fiji using a custom macro to assemble multi-channel hyperstacks from the acquired raw fluorescence images. The resulting hyperstacks were subsequently analyzed in CellProfiler (version 4.2.5) using a custom analysis pipeline (Stirling et al., 2021). Images were grouped according to biological replicate and construct condition using metadata extracted from file names. Analysis was performed on the DAPI, EGFP, rhodamine-phalloidin, and WGA-Alexa Fluor 647 channels. Prior to segmentation, fluorescence intensities were rescaled and images were resized fourfold to improve segmentation accuracy. Nuclear segmentation was performed from the DAPI channel using the StarDist plugin (RunStarDist module) with the pre-trained “Versatile (fluorescent nuclei)” model (Schmidt et al., 2018; Weigert et al., 2020). Segmentation was performed in 2D using a probability threshold of 0.6 and an overlap threshold of 0.1. Nuclei were subsequently filtered based on object size and DAPI intensity to remove debris and low-signal objects. Specifically, only nuclei with an area ≥80 pixels and a mean DAPI intensity ≥0.005 arbitrary units (a.u.) were retained for downstream analysis. Whole-cell segmentation was performed using the Cellpose algorithm (RunCellpose plugin, Cellpose version 2, cyto2 model) (Stringer et al., 2021) based on the WGA membrane signal. The resized DAPI image was supplied as the nuclear reference image during segmentation. Cellpose segmentation was performed without GPU acceleration using a 30-pixel diameter estimation, a flow threshold of 0.4, and a minimum object size threshold of 15 pixels. Objects touching image borders were excluded automatically. Nuclear and cellular objects were subsequently related using CellProfiler parent–child assignment modules to associate individual nuclei with corresponding cells. To restrict analysis to cells expressing EGFP control or EGFP-tagged Ect2 constructs while excluding untransfected cells as well as cells with excessive overexpression, cells were filtered based on mean GFP fluorescence intensity. Only cells within a defined GFP intensity range above background (Minimum value: 0.0025) and below a predefined upper intensity cutoff (Maximum value: 0.4) were retained for downstream analysis.

A broad range of single-cell morphological and fluorescence-based features was extracted for each segmented cell, including cell area, perimeter, mean fluorescence intensities, and shape descriptors. Among these, the form factor (referred to as circularity throughout the manuscript) was used as a quantitative measure of cell shape and calculated according to the following equation:

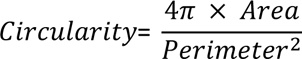

where a value of 1 corresponds to a perfect circle and lower values indicate increasing deviation from circular morphology. F-actin intensity was quantified from the rhodamine-phalloidin channel within the segmented whole-cell masks. Mean fluorescence intensity values were extracted on a per-cell basis using identical segmentation, filtering, and quantification settings across all experimental conditions and biological replicates. For each condition, 60 fields of view were analyzed per experiment. Single-cell measurements were pooled across all analyzed fields and biological replicates for downstream statistical analysis. All downstream data processing, statistical analyses, and graphical representations were performed using GraphPad Prism (version 10.6.1).

#### 3D visualization and dynamic analysis of z-stack time series

3D visualization of fluorescence z-stacks was performed using Fiji. Acquired z-stack images were calibrated according to the acquisition pixel size and z-step spacing prior to rendering. 3D projections were generated using the Fiji 3D Project function with the “Brightest Point” projection method and interpolation enabled. Rotational renderings were generated around the x-axis with a total rotation angle of 360° and rotation increments of 10°. Z-stack slice spacing was set to 0.4 µm. Composite fluorescence channels were merged prior to rendering to preserve multi-channel information in the final projections.

For kymograph analysis, front-view projections generated from the 3D renderings were used. Line regions of interest (ROIs) were manually drawn along the indicated structures on the projected images, and kymographs were generated using the KymographBuilder plugin (https://github.com/fiji/KymographBuilder) in Fiji. Identical rendering and visualization settings were maintained within each experiment.

#### Orthogonal reslice visualization of fluorescence z-stacks

To visualize plasma membrane localization in 3D, fluorescence z-stacks of fixed cells expressing EGFP or EGFP-tagged Ect2 constructs were analyzed in Fiji (ImageJ distribution) using the built-in Dynamic Reslice function. Line regions of interest (ROIs) were manually positioned across the regions of interest, and orthogonal xz views were generated from the original z-stack data to visualize the axial distribution of EGFP fluorescence. Identical visualization settings were applied within each experiment, and representative orthogonal views were exported without nonlinear intensity transformations.

---

**Supplementary Figure 1:**
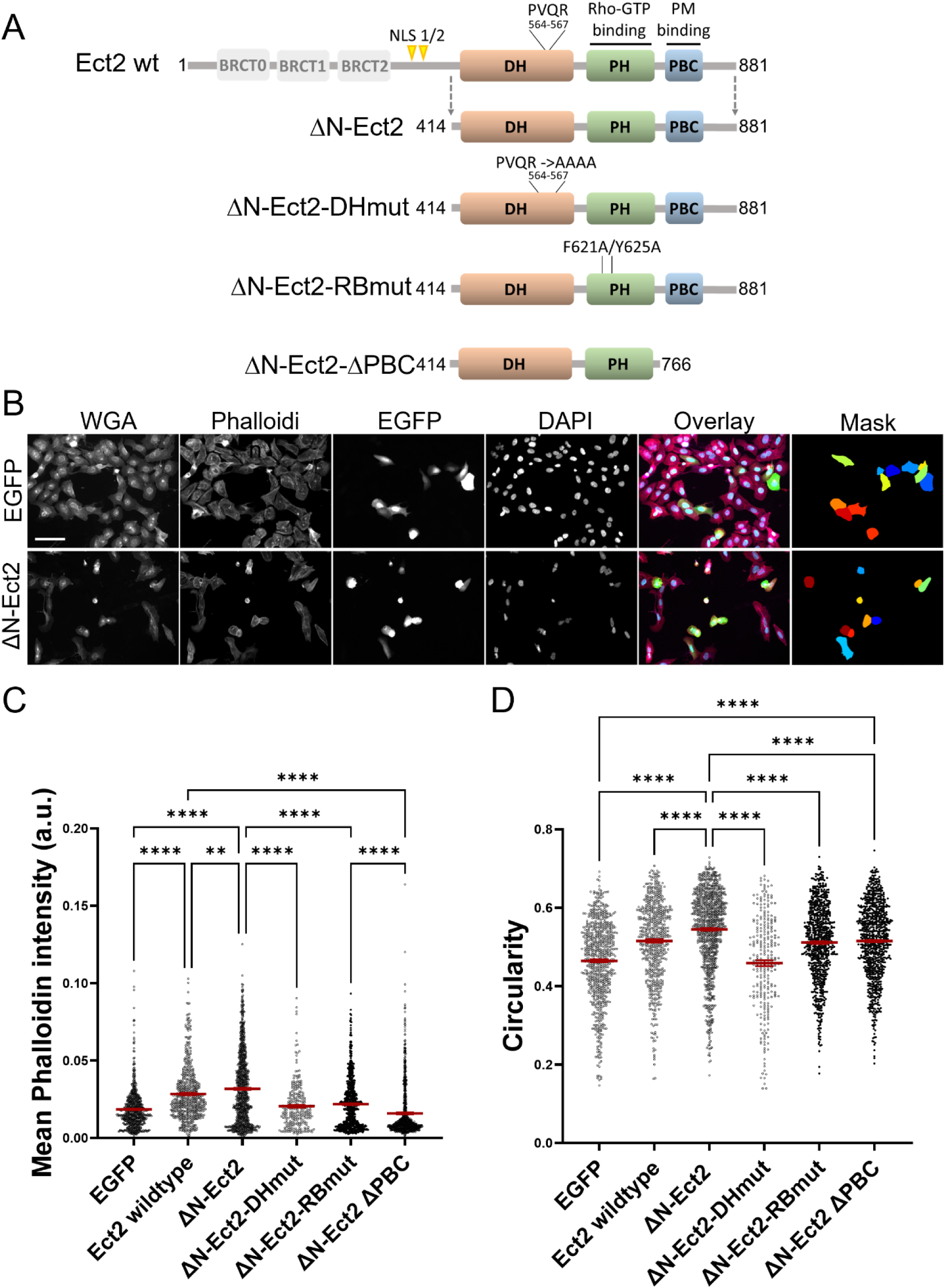
**A)** Schematic domain structures of Ect2 variants. Wild-type Ect2 (Ect2-wt), the N-terminally truncated, constitutively active: ΔN-Ect2 (aa 414-881), catalytically inactive ΔN-Ect2-DHmut, Rho-binding-deficient ΔN-Ect2-RBmut and truncated active variant lacking the C-terminal polybasic cluster for plasma membrane association: ΔN-Ect2-ΔPBC (aa 414-766). **B)** Representative widefield fluorescence images of cells expressing EGFP (control) or EGFP-ΔN-Ect2 stained with WGA (membrane), rhodamine-phalloidin (F-Actin), and DAPI (nuclei). Shown are the individual fluorescence channels, the merged image and the corresponding segmentation mask of detected EGFP-positive (transfected) cells used for quantitative image analysis (described in the methods section). Pseudocolors indicate individual segmented cells. Scale bar = 100 µm. **C-D)** Quantification of mean phalloidin fluorescence intensity and circularity of cells expressing the indicated constructs shown in **(A).** Phalloidin intensity and cell circularity were quantified by automated image analysis as described in the methods section. Data represent n ≥ 276 cells from 4 independent experiments. Statistical analysis was performed using one-way ANOVA followed by Tukey’s post hoc test. Bars indicate mean ± SEM.

**Supplementary Figure 2:**
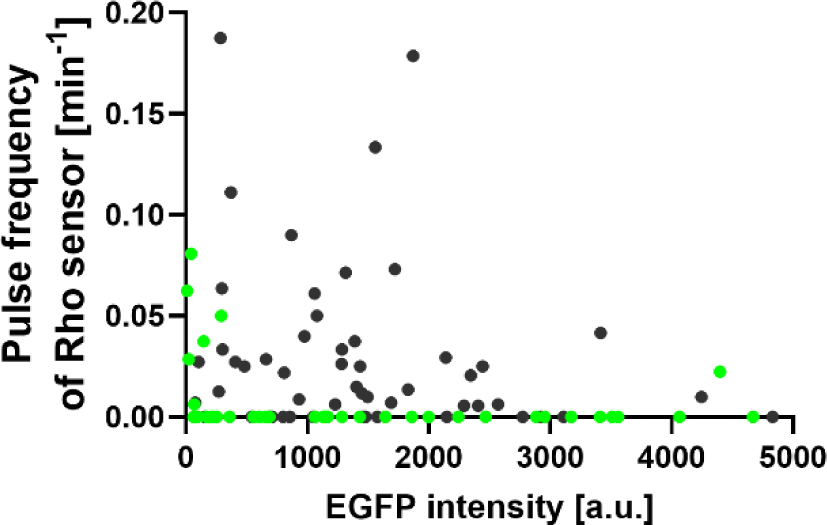
Background-corrected average EGFP intensity in U2OS cells expressing EGFP control (black dots) or EGFP-ΔN-Ect2 (green dots), plotted against the normalized mean pulse frequency of the RBD sensor. Each dot represents a single cell. Data are from n ≥52 cells across three independent experiments.

**Supplementary Figure 3:**
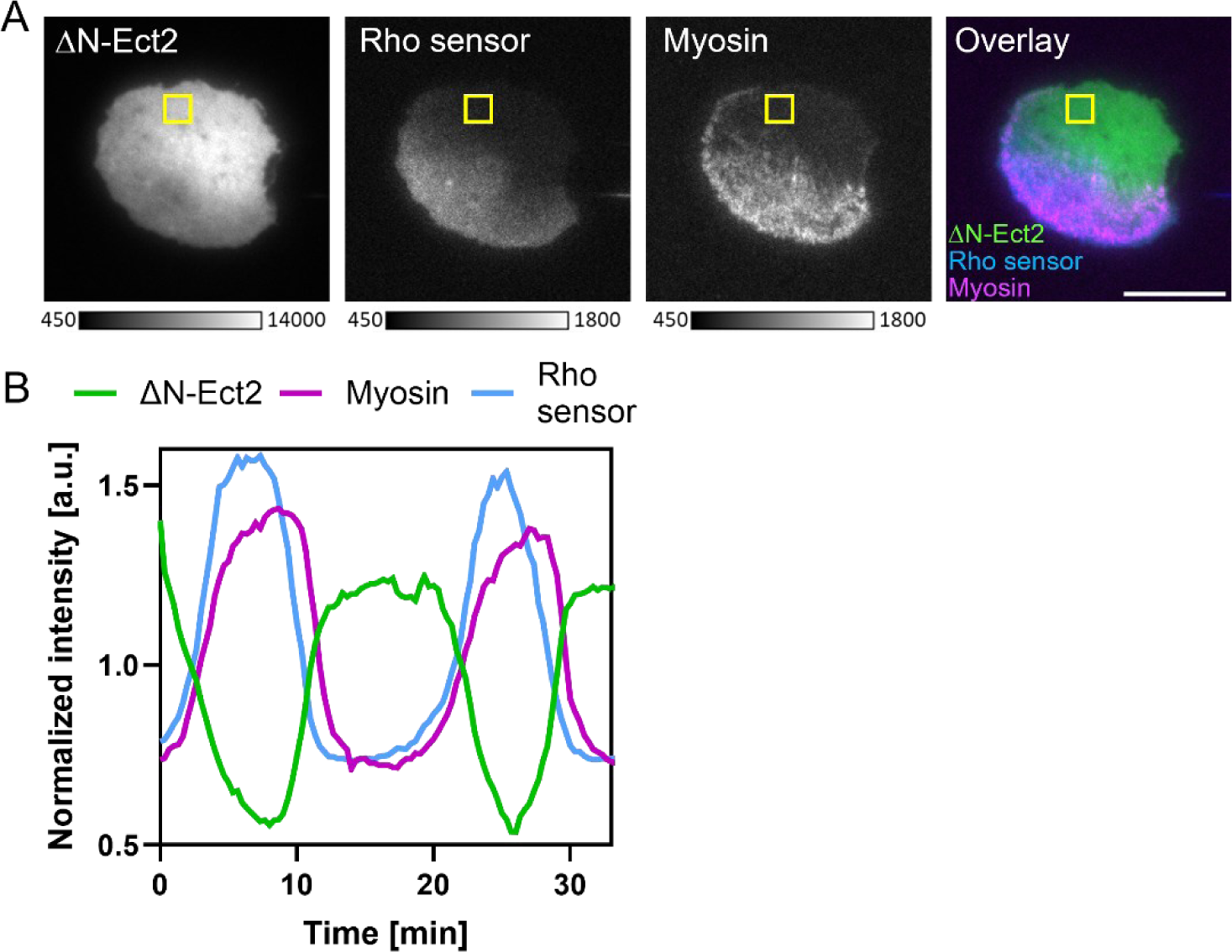
**A)** TIRF images of a representative cell co-expressing EGFP-ΔN-Ect2, a second-generation Rho activity sensor containing two tandem RBD domains delCMV-mCitrine-2xRBD (Nanda et al., 2023) and mCherry-Myosin-IIA. **B)** Signal intensity profiles corresponding to the regions indicated in **(A)**. n=26 cells from 5 experiments. Frame rate: 3 frames/min, scale bar = 20 µm.

## Supplementary Videos

Video 1: **Active, plasma membrane-bound Ect2 suppresses spontaneous pulsatile Rho activity.** TIRF videos of representative cells co-expressing the Rho activity sensor (mCherry-Rhotekin RBD) and either EGFP control, constitutively active Ect2 (EGFP-ΔN-Ect2) or constitutively active GEF-H1 (EGFP-GEF-HI C53R). Two phenotypes induced by active Ect2 are shown: Panel 2: Reduced pulsatory Rho sensor signal, Panel 3: Peripheral enrichment of Rho sensor signal with slow circumferential movement. Frame rate: 3 frames/min. Scale bar = 20 µm.

Video 2: **Active Ect2 suppresses Lbc-GEF stimulated Rho activity pulses.** TIRF time-lapse imaging of cells expressing a Rho activity sensor together with EGFP control (top panels) or constitutively active Ect2 (EGFP-ΔN-Ect2; bottom panels). Left panels show cells prior to nocodazole treatment, whereas right panels show the same cells 45 min after nocodazole addition. Frame rate: 3 frames/min. Scale bar = 20 µm.

Video 3: **Active plasma-membrane bound Ect2 inhibits Myosin-IIA contraction pulses and generates strong peripheral Myosin-IIA enrichment**. TIRF time-lapse imaging of cells expressing mCherry-Myosin-IIA and either EGFP control (left panel) or constitutively active Ect2 (EGFP-ΔN-Ect2; right panels). Two phenotypes induced by active Ect2 are shown. Panel 2: Reduced pulsatory Myosin-IIA signal dynamics, Panel 3: Peripheral enrichment of Myosin-IIA with slow circumferential movement. Frame rate: 3 frames/min, scale bars = 20 µm.

Video 4: **Circumferential cortical Myosin-IIA dynamics are associated with oscillatory cell tilting.** Three-dimensional renderings of spinning disk confocal z-stack time-lapse images. Cells co-express mCherry-Myosin-IIA (magenta) and constitutively active Ect2 (EGFP-ΔN-Ect2) (green). The top panel displays the tilted view highlighting circumferential movement of cortical Myosin-IIa, whereas the bottom panel shows the same cell from a frontal view, demonstrating a periodic rocking motion of the cell. Frame rate: 3 frames/min, scale bar = 5 µm.

Video 5: **Distinct phase relationships between RhoGEFs and Rho activity.** TIRF time-lapse imaging of cells expressing the Rho activity sensor (magenta) together with constitutively active GEF-H1 (EGFP-GEF-H1 C53R, green; left) or constitutively active Ect2 (EGFP-ΔN-Ect2, green; right). Active GEF-H1 induces centrally localized pulses in which GEF-H1 and Rho activity exhibit synchronous dynamics. In contrast, active Ect2 and Rho activity in peripheral waves display anti-phasic dynamics. Frame rate: 3 frames/min, scale bar = 20 µm.

Video 6: **Three-color imaging of Ect2, Rho activity, and Myosin-IIA dynamics. Related to Video 5.** TIRF time-lapse imaging of cells co-expressing constitutively active Ect2 (EGFP-ΔN-Ect2, green), a second-generation Rho activity sensor containing two tandem RBD domains (delCMV-mCitrine-2xRBD, blue, (Nanda et al., 2023), and mCherry-Myosin-IIA (magenta). Frame rate: 3 frames/min, scale bar = 20 µm.

Video 7: **Plasma membrane binding deficient active Ect2 stimulates Rho activity pulses.** TIRF time-lapse imaging of cells expressing the Rho activity sensor (right) together with EGFP control, constitutively active Ect2 (EGFP-ΔN-Ect2), or a constitutively active variant lacking the C-terminal plasma membrane binding region (EGFP-ΔN-Ect2-ΔPBC) (left). Frame rate: 3 frames/min, scale bar = 20 µm.

